# Co-allocation of distinct contextual memories within a wide time window: temporal dynamics and neuronal co-reactivation during retrieval

**DOI:** 10.1101/2025.08.17.670544

**Authors:** Khalid Saidov, Anna Tiunova, Konstantin Anokhin

## Abstract

Recent studies have shown that a shared neuronal ensemble in the hippocampus links distinct contextual memories encoded within a certain time window (specifically, 5 hours, 2 and 7 days). Here we explored the temporal dynamics of two contextual (neutral and aversive) memories linking and analysed neuronal ensembles in the hippocampus, amygdala and different cortical regions reactivated during retrieval. Firstly, we have found that memories integrated across different time-points including several hours, days and weeks but not if learning phases was separated by short-term and very long-term time intervals. Secondly, we have demonstrated a higher neuronal co-reactivation in the hippocampus and amygdala during retrieval in case of memories integration that supports the hypothesis that shared neuronal ensembles link distinct memories. Finally, we have elicited that proportions of reactivated neuronal ensembles in these brain regions are greater in case of contextual memories integration.

## Main text

Memory may be defined as an internal representation of experience that can be recalled at later time (Dudai, 2007). Such internal representation is thought to be encoded by long-lasting physical changes in neuronal populations that form a memory trace called a memory engram (Dudai 2004; Josselyn et al., 2015). One of the key questions is how neurons are chosen to become recruited into a given memory engram. Previous studies have shown that the transcription factor cyclic adenosine 3′,5′-monophosphate response element binding protein (CREB) plays a crucial role in selection of neurons to become a part of memory engram (Han et al., 2007). For instance, neurons in the lateral amygdala with the relatively high level of CREB expression during learning are preferentially recruited into a fear memory engram and are necessary for the subsequent recall (Han et al., 2007). An artificial reactivation of these neurons was sufficient to recall fear memory without presentation of an external retrieval cue and induced a reconsolidation-like reorganization process (Kim et al., 2013). The crucial role of CREB has been shown not only in the lateral amygdala but also in the insular cortex and hippocampus (Sano et al., 2014). This process of preferential recruiting activated neurons in a memory trace was called memory allocation (Rogerson et al., 2014; Frankland and Josselyn, 2015; Park et al., 2016).

The molecular mechanism of memory allocation in certain neuronal ensembles can be related to the fact that neurons with a high level of CREB expression are more excitable than their neighbours and showed greater synaptic strength following training (Impey et al., 1996; Dong et al, 2006; Zhou et al., 2009; Sano et al., 2014). Increase in the neuronal excitability enhances likelihood that these neurons will be involved into a memory trace (Silva et al., 2009; Yiu et al., 2014). In other words, allocation of memories in the brain is mediated by electrophysiological excitability of neurons: cells with the higher excitability preferentially involve in encoding, storage, and retrieval of these memories.

Remarkably, that the memory allocation hypothesis suggests that CREB activation during learning can induce temporal increase in excitability of neurons involved into a given memory engram that biases the storage of subsequent memories to the same neuronal population that encoded the first memory. This striking prediction supposes that a shared neural ensemble can link distinct episodic memories encoded close in time (Rogerson et al. 2014). Indeed, the recent experimental evidence have demonstrated that two episodic memories that encoded within a few hours can be integrated (Cai et al., 2016; Rashid et al., 2016). This phenomenon was called memory co-allocation (Sehgal et al., 2018). In particular, a shared neuronal population in the CA1 region of hippocampus links distinct contextual memories encoded within 5 hours but not 7 days (Cai et al., 2016). Another study has shown that two fear memories to different sounds were linked when these events occurred closely in time (from 1,5 to 6 hours, but not from 18 to 24 hours) (Rashid et al., 2016). In addition, the possibility to integrate two episodic memories by repetitive co-retrieval has been demonstrated. Thus, the presentation of conditioned stimulus used for the one task can trigger the conditioned response of the other task (Yokose et al., 2017). Such co-retrieval reorganizes two memories traces to form a shared neuronal ensemble that links them (Yokose et al., 2017). Moreover, it was demonstrated that in mice, a strong aversive experience drives offline ensemble reactivation of not only the recent aversive memory but also a neutral memory formed 2 days before, linking fear of the recent aversive memory to the previous neutral memory. Fear specifically links retrospectively, but not prospectively, to neutral memories across days (Zaki et al., 2024).

At the same time, the temporal dynamics of memory co-allocation across different time intervals (from short to very long) and the neuronal activation in a broad number of traditionally associated with memory brain regions during linked memories retrieval remain the subject of further studies. To explore this temporal dynamics, we used the behavioural paradigm partially based on the contextual fear conditioning in mice.

All experimental procedures were carried out in accordance with the Directive 2010/63/EU of the European Parliament and of the Council of the European Union issued September 22, 2010, on the protection of animals used for scientific purposes (Section 27) and in line with Order №267 Ministry of Healthcare of the Russian Federation (19.06.2003) and Decision by the Local Ethical Committee of Biomedical Research by National Research Centre Kurchatov Institute (Protocol №1, 09.07.2015). Adult C57BL/6J mice (10-14 weeks old) were housed in groups of 3-4 and maintained on a 12:12 hours light/dark cycle with light from 9 a.m. to 9 p.m. and ad libitum access to food and water.

Separate groups of mice (n=15 in each group) were allowed to explore a neutral context A for 10 minutes and after 5 minutes, 30 minutes, 2 hours, 5 hours, 1, 3, 7, 30 or 120 days were placed to an aversive context B, where they received a foot shock (Figure 1A). The protocol of training in the aversive context was the following: each mouse explored the fear conditioning chamber for 3 minutes and then received three 2 second foot shocks of amplitude 1.5 mA with an intershock interval 30 seconds. Then, 1 minute after the final shock, the mice were removed and returned to the vivarium. 72 hours after the training in the aversive context all mice were tested for 5 minutes in the contexts A, B, and the completely novel context C which was not exposed earlier. The testing sequence in all the contexts was counterbalanced (i.e. 1/3 of mice from each group were tested in ABC, BAC, CAB on the following three days, respectively) with intervals 24 hours between testing days. All testing was done in Med Associates chambers (MED Associates Inc.). Learning and tests in each context were performed in different experimental rooms. Behavioural data were processed using the Med Associates software for measuring freezing. Design of all the used contexts see in the Supplemental Material (Figure S5).

**Figure 1.**
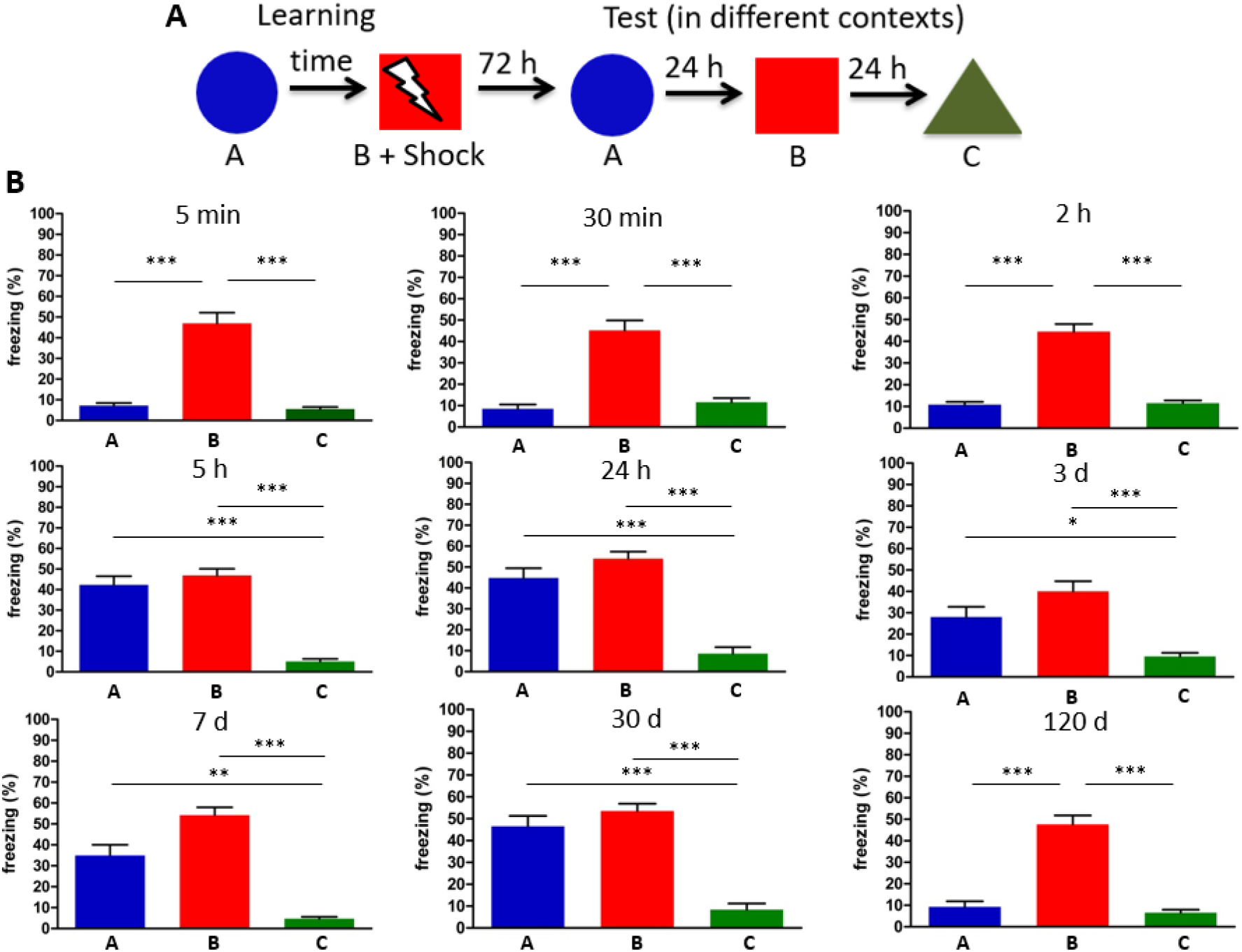
Two distinct contextual memories (the neutral and aversive) can be linked if they encoded within intermediate and long-term time intervals. (A) The behavioural paradigm in which the aversive memory associated with a foot shock in context B specifically transferred to the memory in the neutral context A. (B) Mice (n=15 in each group) froze both in the context A (initially neutral) and B (associated with the foot shock) but not in the completely novel context C if they have been trained in the contexts A and B within different time intervals (from 5 hours until 1 month). P < 0.05 (Dunn’s Multiple Comparison test). Bars indicate mean ± s.e.m.

We have found that a delay from 5 hours including to 30 days between trainings resulted in significantly higher freezing during the test in the neutral context along with the aversive context but not in the completely novel context (P < 0.05), indicating the transfer of the fear memory to the neutral memory (Figure 1B). In contrast, this did not occur if animals were trained in two contexts within 5 minutes, 30 minutes, 2 hours and 120 days (P < 0.05) (Figure 1B). In other experiment we checked if this effect depends on the specific design of context A used in our experiments. We trained a separate group of mice (n=15) in the same paradigm within 24 hours but reversed the neutral and previously completely novel contexts (i.e. used a design of the completely novel context C as the neutral context) then trained in the aversive context B and tested in contexts C, B and the completely novel context A (Figure S1A). The testing was performed according to the same paradigm as in the previous experiments. We have observed that changing the neutral context design did not influence on the aversive memory transfer to the neutral context and this effect did not generalize during the test in the completely novel context (P < 0.05) (Figure S1A). In another experiment, we first trained a separate group of mice (n=15) in the aversive context B, then 24 hours after exposed the neutral context A, tested in the same manner and have found that training initially in the aversive context did not lead to the aversive memory transfer to the neutral context (P < 0.05) (Figure S1B).

Next, we used the two-colour c-fos catFISH method (Saidov and Anokhin, 2018) to visualize and compare neuronal ensembles in the hippocampus, amygdala and different cortical regions activated during retrieval of neutral and aversive memories encoded within time intervals that led or not to the behavioural linking. To this aim we trained two groups of mice (each n=4) in the same behavioural paradigm within 24 hours or 5 minutes. Animals were trained in the contexts A and B then 72 hours after were tested for 5 minutes initially in the completely novel context C then 24 hours after were tested on the same day for 5 minutes in the contexts A and B within 20 minutes before sacrifice (Figures 2A, S2). This approach allowed us to tag neuronal ensembles reactivated in the neutral and aversive contexts. We used the previously described protocol of the two-colour c-fos catFISH (Saidov et al., 2019) to visualize and count cells containing the cytoplasmic (Cyto^+^), nuclear (Nu^+^) and both the nuclear and cytoplasmic (Nu^+^ + Cyto^+^) mRNA (i.e. ensembles reactivated in the contexts A, B and co-reactivated in both contexts, respectively) (Figure S6). We selected three coronal brain slices from each brain and analysed c-fos mRNA positive cells as previously described (Saidov et al., 2019). The formulas for cell counting are described in the Supplemental Material (table 1).

**Table 1.**
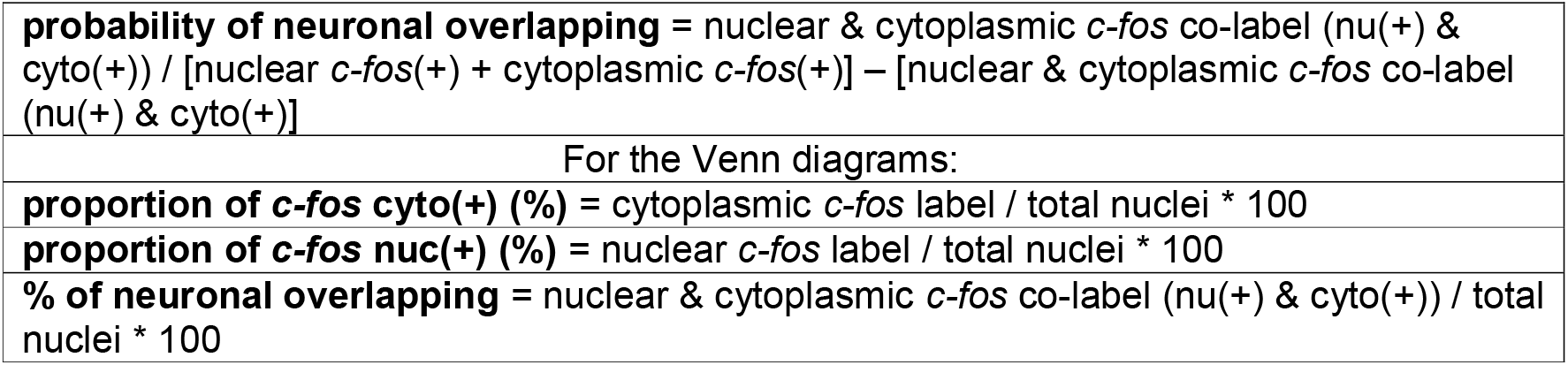
The formulas for cell counting.

**Figure 2.**
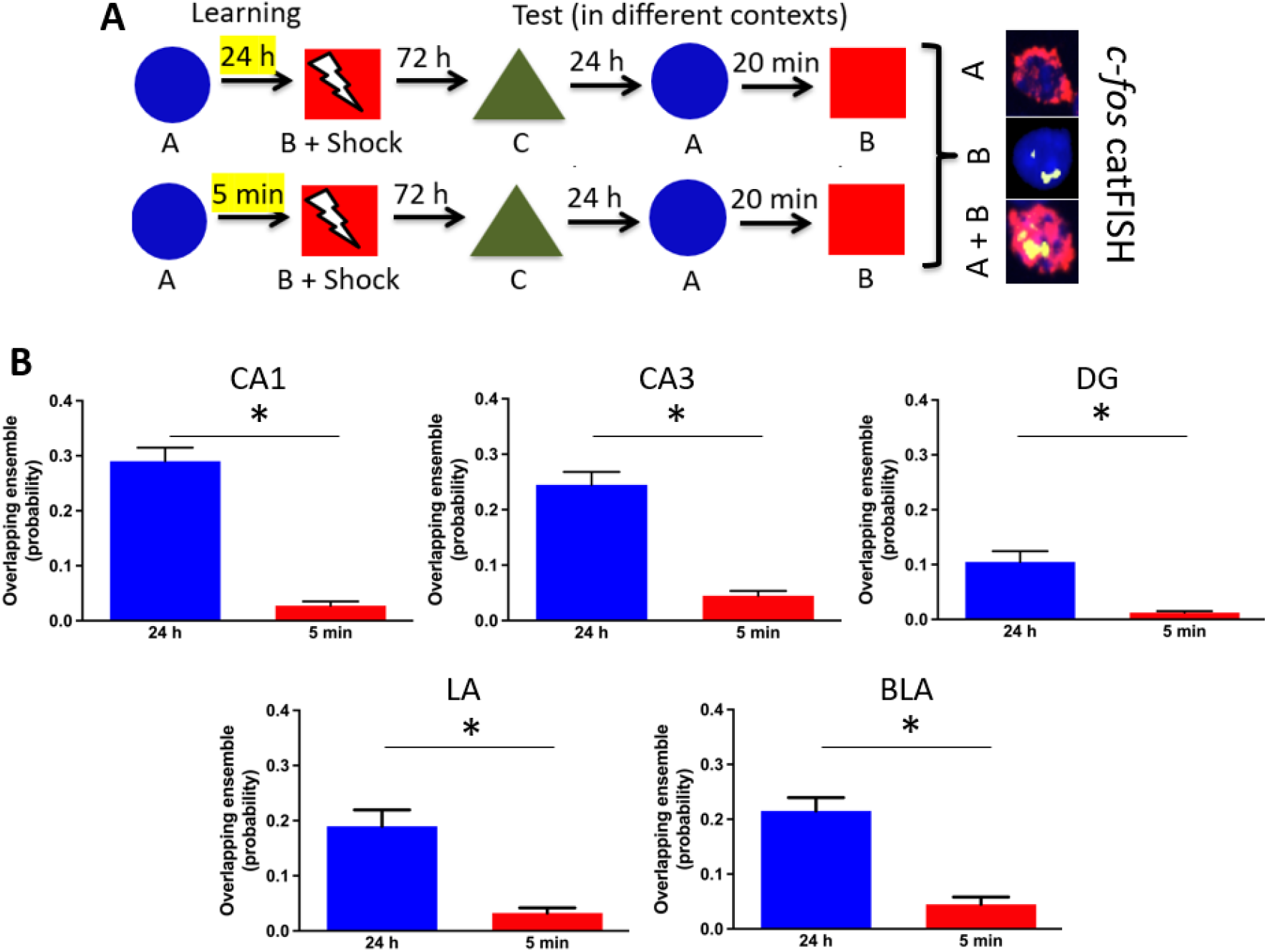
Shared neuronal ensembles in the hippocampus and amygdala co-activated during retrieval in contexts A and B in case of aversive memory transfer. (A) Two groups of mice (n=4 in each group) were trained in the contexts A and B within 24 hours or 5 minutes. After retrieval and sacrifice, brains were stained according to the two-colour c-fos catFISH protocol to analyse neuronal populations reactivated in contexts A, B and co-reactivation in both contexts. (B) Co-reactivation of neuronal populations in the cornu ammonis 1, 3 (CA1, CA3), dentate gyrus (DG) of hippocampus, lateral (LA) and basolateral (BLA) amygdala after retrieval in contexts A and B is significantly higher if mice have been learnt in contexts A and B within 24 hours but not 5 min. P < 0.05 (Mann-Whitney test). Bars indicate mean ± s.e.m.

An overlap of ensembles activated in contexts A and B was significantly higher in the hippocampus, specifically, in the cornu ammonis 1, 3 (CA1, CA3), dentate gyrus (DG) and lateral and basolateral amygdala (LA, BLA) if neutral and aversive memories encoded within 24 hours but not 5 minutes (P < 0.05) (Figure 2B). In contrast, there was not significant difference between co-activated in two contexts neuronal populations in cortical areas (Figure S3). We analysed the following regions of cortex: frontal associative (FrA), prelimbic (PrL), infralimbic (IL), cingular 1 and 2 (Cg2, Cg2), retrosplenial (RSD), lateral parietal associative (LPtA), medial parietal associative (MPtA).

We also analysed the proportion of ensembles reactivated in the contexts A and B and have found that the size of neuronal populations in the hippocampus and amygdala reactivated in contexts A and B are greater in case of neutral and aversive contextual memories linking. Namely, the ensemble reactivated during the test in the context A is bigger in the CA1 if mice have been learnt within 24 hours but not 5 minutes (P < 0.05) (Figure 3). In the meantime, the number of reactivated populations both in the LA and BLA is higher after retrieval in contexts A and B (P < 0.05) (Figure 3). In contrast, cortical neuronal ensembles reactivated in contexts A and B after learning within 24 hours or 5 minutes is relatively similar (Figure S4). Thereby, the transfer of the aversive memory to the neutral memory led to increase of cell amount reactivated during retrieval.

**Figure 3.**
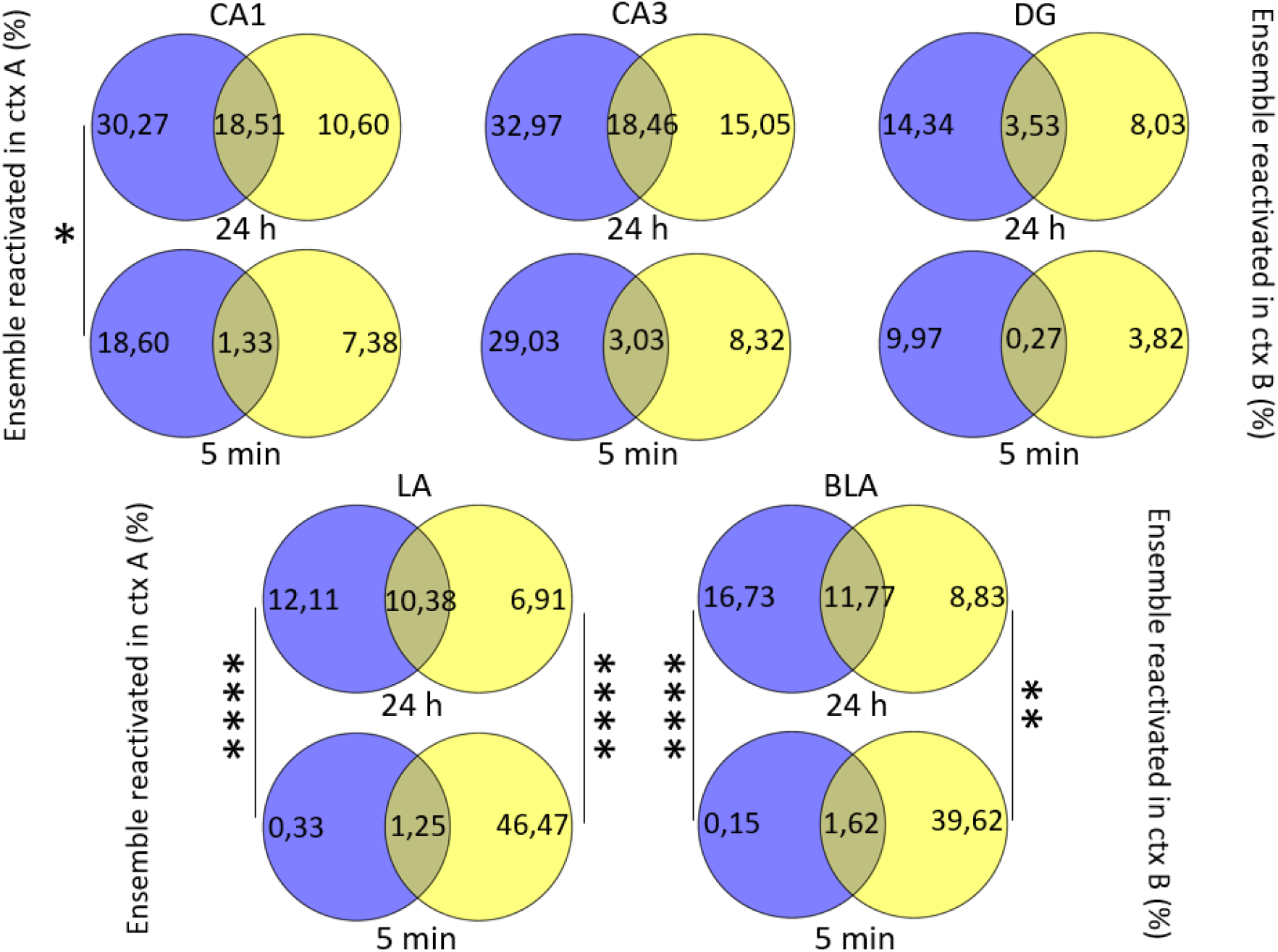
The proportion of neuronal ensembles of the hippocampus and amygdala reactivated in contexts A and B is bigger in case of the neutral and aversive contextual memories linking. The Venn diagrams show that the number of cells reactivated during the test in the context A is higher in CA1 if mice have been learnt within 24 hours but not 5 minutes. In LA and BLA the size of reactivated populations is bigger both after retrieval in the contexts A and B. P < 0.05 (Mann-Whitney test). The overlapping field on the Venn diagrams reflects percent of co-activated cells.

Results of the present study have demonstrated that two distinct contextual memories (in case of our study the neutral and aversive) can link in mice within a wide time window which span from several hours to days and weeks. We have discovered distinct temporal boundaries in time, when two contextual memories (neutral and aversive) can be linked or not and have found that they integrate (i.e. co-allocate) within not only the intermediate (several hours) but also within different long-term time intervals (days and weeks). Although the neutral and aversive contexts in which animals have been learnt had some similar features, the aversive memory transfer to the neutral memory is specific because did not generalize if animals explore the completely novel context during the test. This result is consistent with the previous findings that reported the possibility for co-allocation of memories encoded close in time (e.g. within several hours) (Cai et al, 2016, Rashid et al., 2016) and 2 or 7 days (Zaki et al., 2024). Recent finding has shown impossibility of the prospective memory linking (i.e. if the aversive experience precedes the exploration of the neutral context). In our study we also did not observe the prospective memory linking using the similar behavioural paradigm but other time intervals between learning in two contexts and between learning/memories recall (24 and 72 hours in our study versus 48 and 24 hours in the previous research respectively). This consistent with the classical notion that the associative memory can not be formed if the unconditioned stimuli (US) precedes the conditioned stimuli (CS) (Pavlov, 1927).

We have also found that the neuronal co-activation during retrieval in the hippocampus and amygdala is higher if distinct contextual memories encoded within 24 hours but not 5 minutes. This result suggests that shared neuronal ensembles reactivate during linked memories retrieval and call up with the previous findings that have demonstrated a neuronal overlap in the CA1 during the two memories integration phase (Cai et al., 2016, Zaki et al., 2024). Surprisingly, that there was no significant distinction of the neuronal co-activation between groups 24 hours and 5 minutes in different cortical areas traditionally associated with memory acquisition and recall.

Finally, our study has shown that memories linking accompanies with the increased proportion of reactivated neuronal ensembles both in the hippocampus (CA1) and amygdala (LA, BLA).

One of the opened questions for further studies is whether and to what extent memory co-allocation across the long-term time window determines by reactivation of neuronal representations for the first memory during the second learning episode or/and offline reactivations after learning within memory consolidation.

Taken together, our findings expand growing knowledge about possibilities of memories linking across time and underlying neural mechanisms.

## Acknowledgements

The study was supported by the Russian Foundation for Basic Research (RFBR) Grant No. 16-04-01545 A

## Generative AI statement

The authors declare that no Generative AI was used in the creation of this manuscript.

## Supplemental_Materials

**Figure S1.**
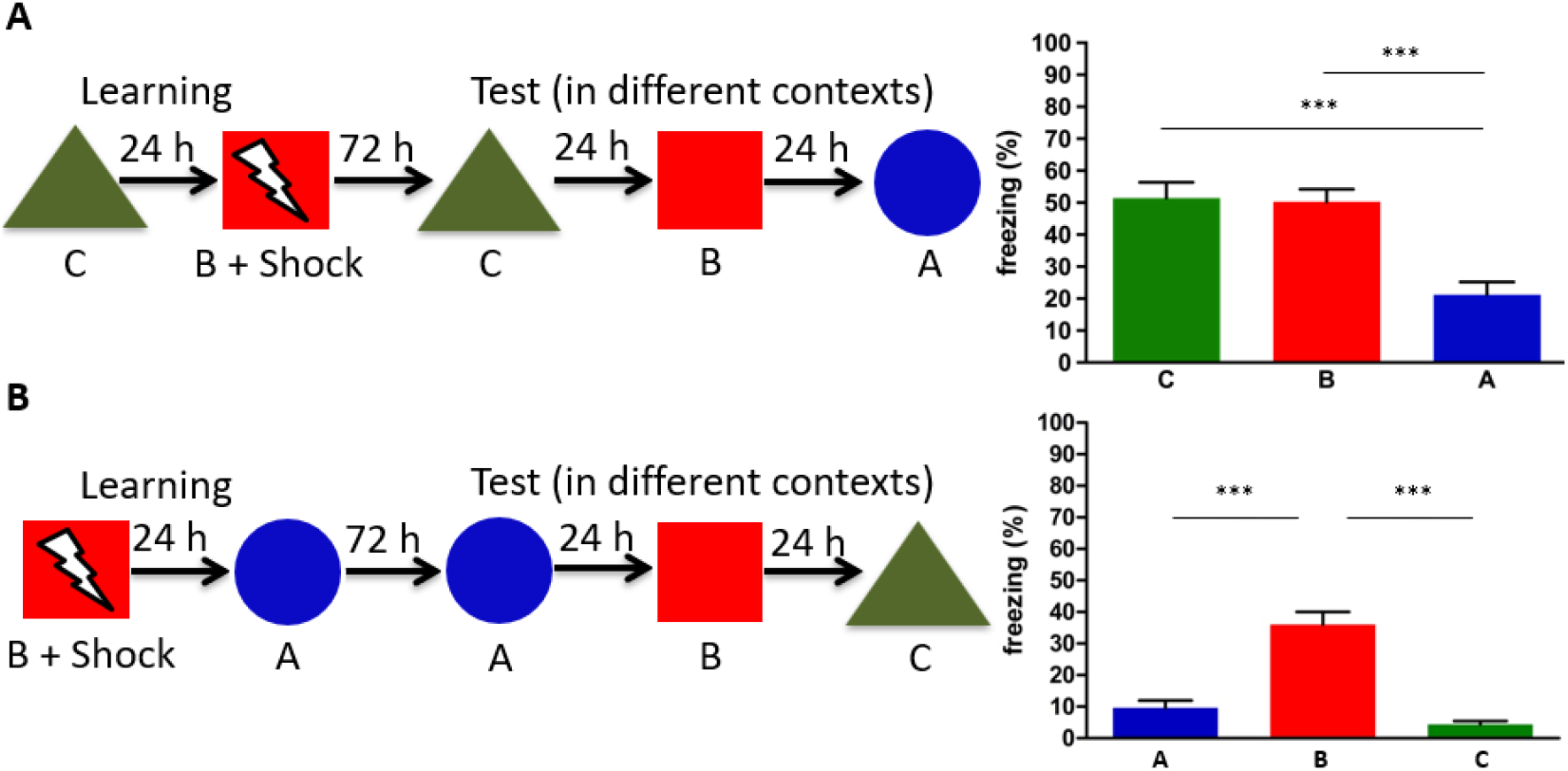
(A) Linking the neutral and aversive contextual memories did not depend on the specific design of initially neutral context. The separate group of mice (n=15) was learnt within 24 hours in the same behavioural paradigm, but previously neutral and novel contexts A and C were reversed. Changing of contexts did not influence on the aversive memory transfer and this effect did not generalize during the test in the completely novel context. (B) Training initially in the aversive context did not lead to two distinct memories integration. The separate group of animals (n=15) was initially trained in the context B associated with the foot shock and then 24 hours after in the neutral context A that did not lead during the test to the aversive memory transfer. P < 0.05 (Dunn’s Multiple Comparison test). Bars indicate mean ± s.e.m.

**Figure S2.**
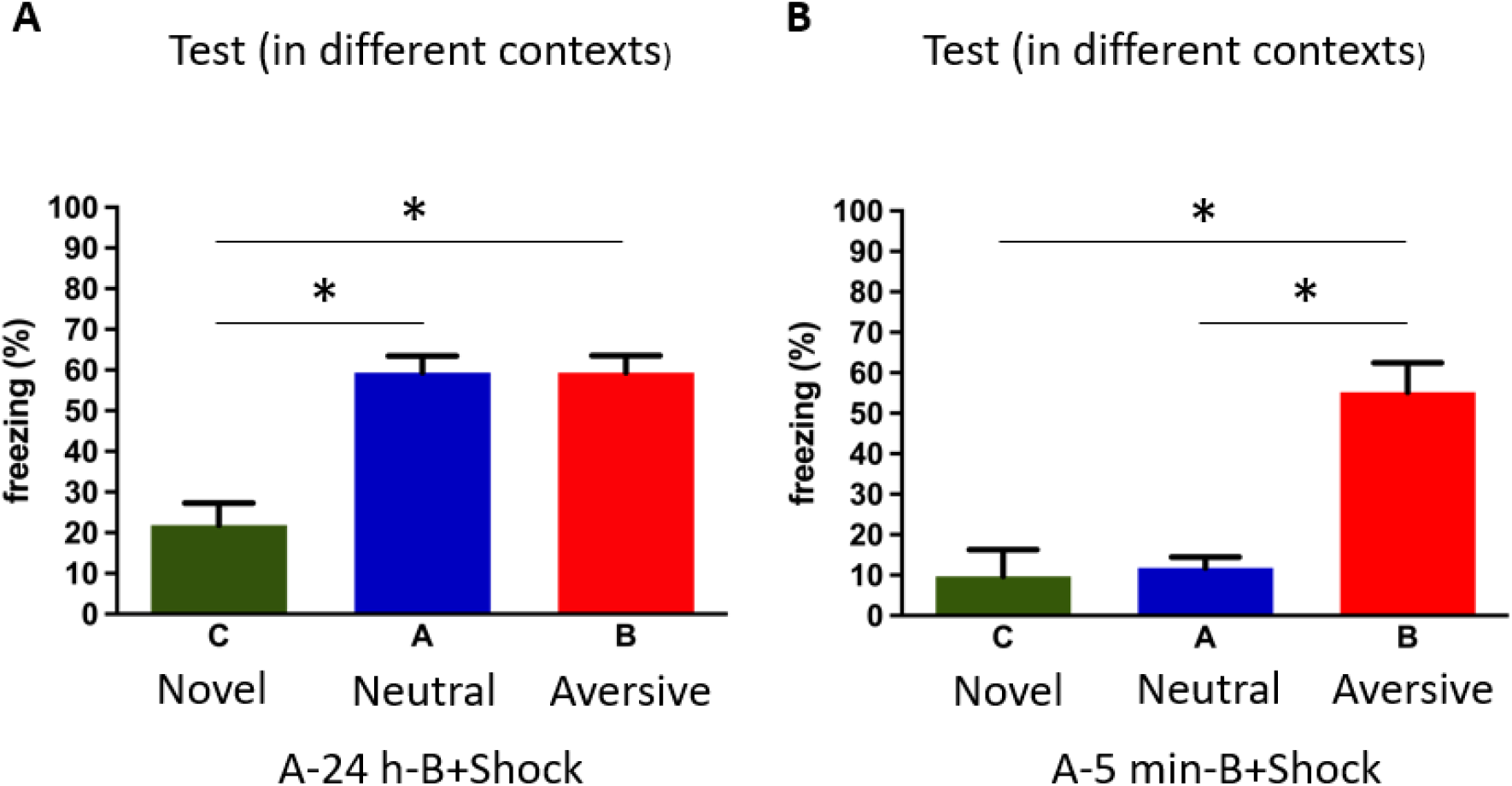
(A, B) Results of the behavioural test in different contexts before animals sacrifice and the two-colour c-fos catFISH staining (related to figure 2A). P < 0.05 (Dunn’s Multiple Comparison test). Bars indicate mean ± s.e.m.

**Figure S3.**
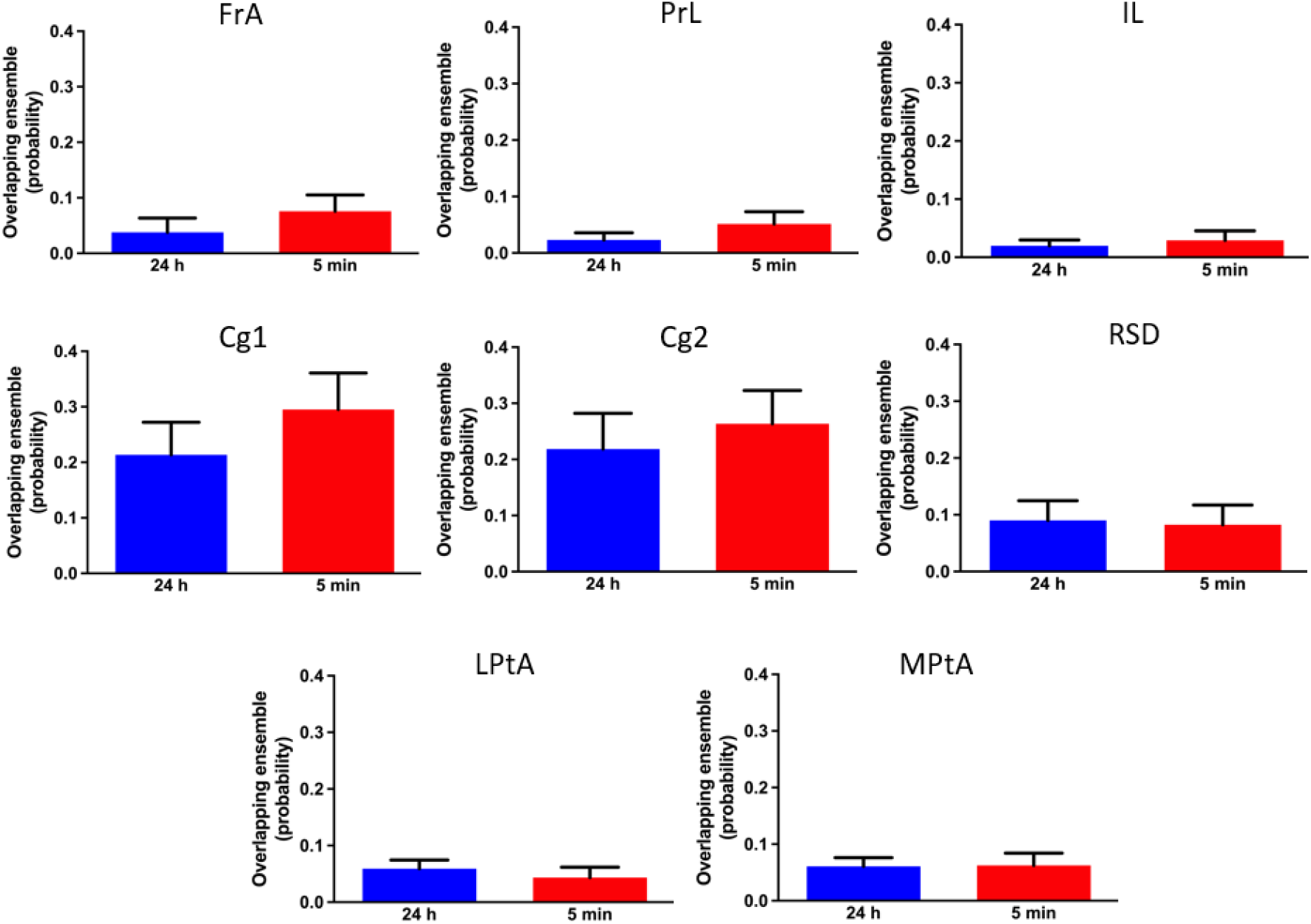
Two distinct contextual memories linking was not accompanied with a higher neuronal overlap in different cortical areas. The amount of co-activated cells in different regions of cortex was not significantly higher after retrieval in contexts A and B in case of aversive memory transfer. Abbreviations: FrA-frontal associative, PrL-prelimbic, IL-infralimbic, Cg1, Cg2-cingular 1 and 2, RSD-retrosplenial, LPtA-lateral parietal associative, MPtA-medial parietal associative. P < 0.05 (Mann-Whitney test). Bars indicate mean ± s.e.m.

**Figure S4.**
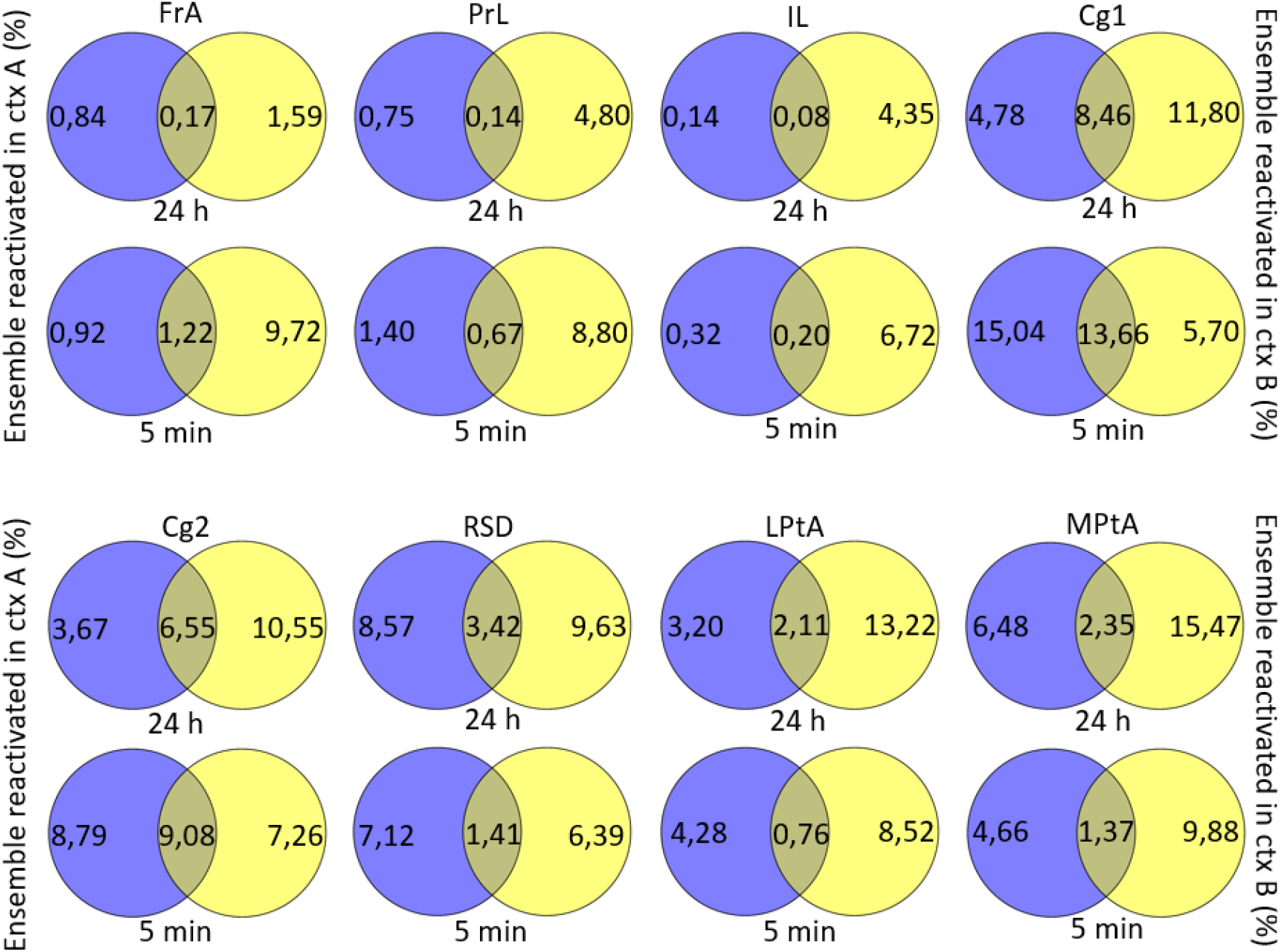
The proportion of different cortical neuronal ensembles reactivated in contexts A and B after learning within 24 hours or 5 minutes is similar. The overlapping field on the Venn diagrams reflects percent of co-activated cells.

**Figure S5.**
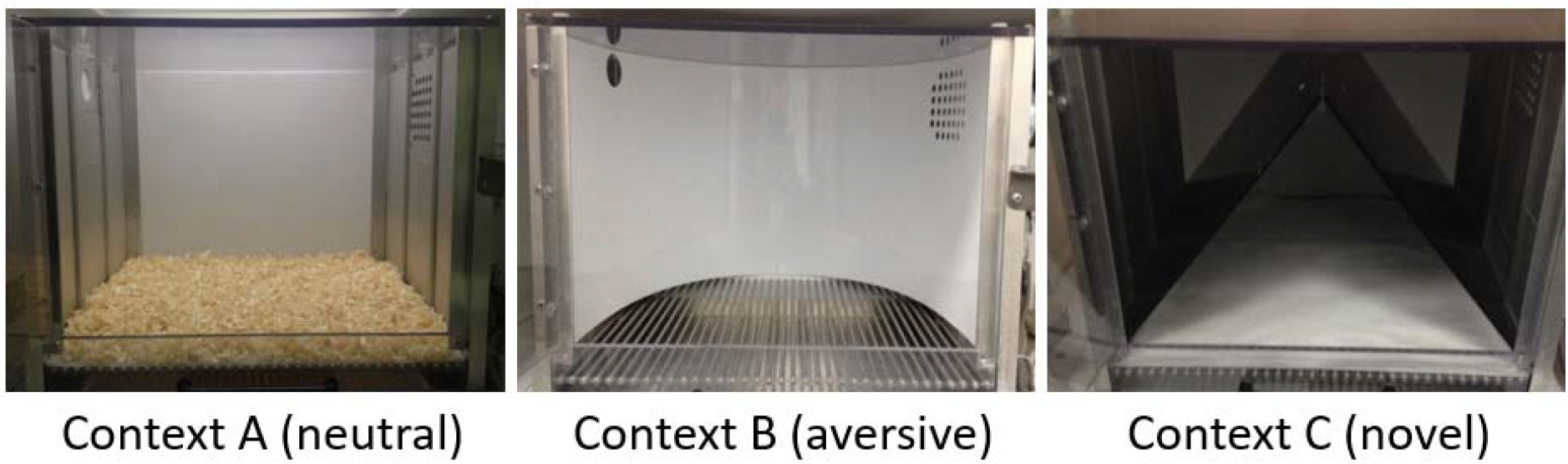
Designs of all the contexts used in experiments.

**Figure S6.**
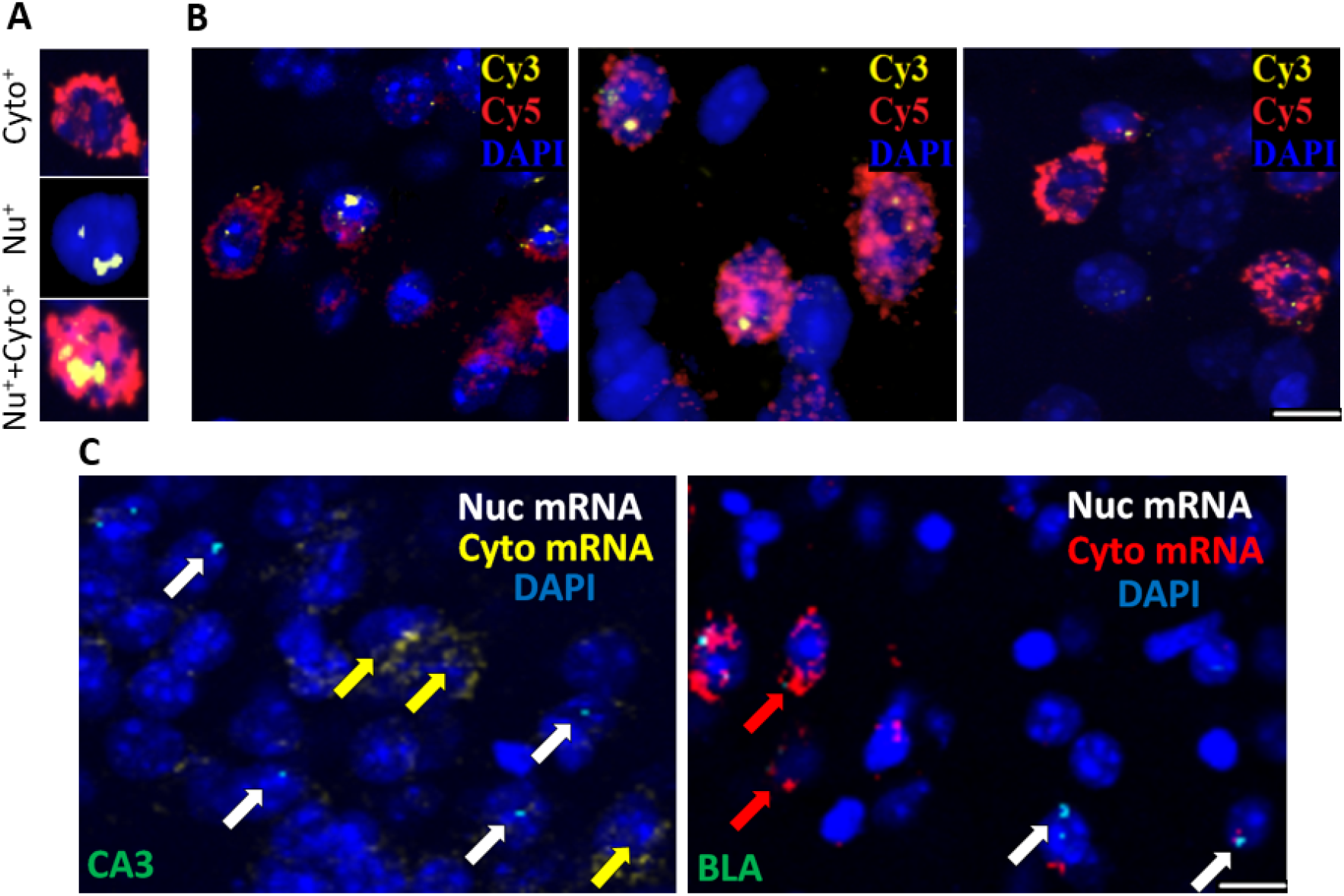
(A) Examples of cells stained by two-colour c-fos catFISH (Abbreviations: Cyto^+^-cytoplasmic, Nu^+^-nuclear, Nu^+^+Cyto^+^-both nuclear and cytoplasmic mRNA). (B) Images showing cells stained by two-colour c-fos catFISH (nuclear, cytoplasmic mRNA and cell bodies stained with dyes Cy3, Cy5 and DAPI respectively). (C) Two-colour c-fos catFISH visualization of ensembles in the CA3 and BLA activated during retrieval in the used behavioural paradigm. Cells that containing cytoplasmic, nuclear and both markers were activated in the contexts A, B or both, correspondingly. Scale bar: 20 μm.

